# Concomitant ablation of SOS1 and SOS2 triggers a lethal phenotype involving compromised intestinal integrity and widespread septicemia

**DOI:** 10.64898/2026.03.20.712875

**Authors:** Andrea Olarte-San Juan, Pablo Rodriguez-Ramos, Alba Diaz-Alguilera, Nuria Calzada, Carmela Gómez, Rocío Fuentes-Mateos, Alberto Fernandez-Medarde, Rubén Nogueiras, David Díaz, Eugenio Santos, Rósula García-Navas

**Author notes:** **Corresponding authors:** Eugenio Santos; Rosula García-Navas.

## Abstract

The RAS guanine nucleotide exchange factors Son of Sevenless 1 and 2 (SOS1 and SOS2) are key regulators of RAS signaling pathways controlling cellular proliferation, differentiation, and survival processes that are essential for correct tissue homeostasis. While mice lacking both SOS1 and SOS2 die precipitously, we demonstrate herein that the combined genetic ablation of SOS1 and SOS2 triggers spontaneous, gut-derived, lethal bacteremia. Double-knockout (DKO) SOS1/2 mice exhibit extensive intestinal tissue damage, massive bacterial leakage out of the gut, and rapid progression to multi-organ failure and death. At the cellular level, loss of both SOS1 and SOS2 leads to profound immune cell depletion and a marked reduction in intestinal stem cell abundance and proliferative capacity, which is accompanied by severe disruption of intestinal architecture and increased epithelial permeability, indicating a breakdown of gut barrier integrity. Notably, therapeutic interventions aimed at enhancing cellular stemness significantly improve survival in SOS1/2 DKO mice, restoring intestinal proliferation and tissue organization. Collectively, our findings identify SOS1 and SOS2 as critical regulators of intestinal homeostasis and regenerative capacity during systemic infection and reveal stemness reinforcement as a potential strategy to overcome lethal susceptibility to sepsis.

## INTRODUCTION

RAS GTPases, originally identified for their oncogenic potential, constitute the most frequently mutated family of oncogenes and drive a wide range of malignancies. These proteins act as molecular switches, cycling between an active GTP-bound state and an inactive GDP-bound state, a process tightly regulated by guanine nucleotide exchange factors (GEFs) and GTPase-activating proteins (GAPs) (1–5).

Among the major RAS-GEF families, SOS proteins are the most ubiquitously expressed and functionally relevant mediators of upstream RAS activation(6). Despite their high structural and sequence homologies, *in vivo* animal models have revealed non-redundant roles for the SOS1 and SOS2 GEFs. *Sos1*-null homozygous mice die during midgestation(7,8), whereas SOS2-deficient animals remain viable(9). To overcome the embryonic lethality associated with SOS1 deletion, we generated conditional, TMX-inducible *Sos1* knockout mouse models, allowing the parallel investigation of SOS1 and SOS2 function in adult animals and tissues. This strategy enabled the establishment of four distinct SOS genotypes (WT, SOS1 KO, SOS2 KO, and SOS1/2 DKO) for comparative analyses. While adult SOS1/2 DKO mice exhibited rapid and severe lethality, both SOS1 KO and SOS2 KO animals remained viable(10).

The exact, underlying cause of death in SOS1/2 DKO mice remains unresolved, as extensive analyses failed to identify a single physiological defect sufficient to account for their lethality(10). Given the central role of SOS GEF proteins in RAS-MAPK pathways regulating cellular proliferation and differentiation, as well as organismal development in different physiological and pathological contexts (6,11–21), here we focused on testing the intestine as a potential site of dysfunction through detailed studies of intestinal homeostasis and functionality, along with analysis of its epithelial renewal and maintenance.

Although direct evidence linking SOS1 and SOS2 to small intestinal function is limited, their established role in intracellular signaling suggests a potential contribution to intestinal physiology. Intestinal epithelial cells (IECs) undergo continuous self-renewal, sustained by a self-renewing intestinal stem cell (ISC) niche every 3-4 days, a process critical for maintaining epithelial barrier integrity(22). Consistently, dysregulated RAS signaling, including aberrant RAS activation, has been implicated in multiple intestinal pathologies, most notably colorectal cancer(23,24).

While the specific functions of SOS proteins in the intestine remain poorly characterized, the mitogen-activated protein kinase (MAPK) pathway is known to play a fundamental role in intestinal homeostasis. Individual components of the MAPK cascade exert distinct effects. For example, Shp2/MAPK signaling regulates goblet and Paneth cell fate decisions and influences lineage allocation and Paneth cell maturation(25). Reduced MAPK activity results in the accumulation of immature Paneth cells(26). Moreover, loss of ERK1 and ERK2 in intestinal epithelial cells impairs nutrient absorption, epithelial migration, and secretory lineage differentiation(27). Recent work has further highlighted the essential role of ERK1/2 in intestinal development through modulation of Wnt signaling via both cell-autonomous and non-cell-autonomous mechanisms(28).

Intestinal homeostasis depends on a finely tuned balance of proliferative signals governed by the RTK-SOS-RAS axis. Consistent with this notion, it has been reported that *Act1* functions as a negative regulator of FGF2-induced ERK signaling by competing with SOS1 for GRB2 binding. During intestinal injury, however, cooperative interactions between regulatory T cells and Th17 cells promote epithelial repair via IL-17 overexpression, which disrupts Act1-GRB2 association(29). In addition, SOS1-driven proliferative signaling in epithelial cells is counterbalanced by the activity of another RAS-GEF, GRP1, contributing to the regulation of normal intestinal epithelial homeostasis and colorectal cancer progression(6,30).

Given the apparent functional interplay between SOS1 and SOS2 in intestinal maintenance, we sought to investigate their combined role in this tissue to try to elucidate the mechanisms underlying the lethality observed in SOS1/2 DKO mice.

## MATERIAL AND METHODS

### Generation of tamoxifen-inducible, *Sos1*-null mice

To obtain *Sos1*-null mice, a mouse strain harboring a floxed version of *Sos1* with exon 10 flanked by LoxP sites (*Sos1^fl/fl^*) was crossed with a mouse expressing tamoxifen (TMX)-inducible Cre recombinase to generate *Sos1^fl-Cre^*/ *Sos1^fl-Cre^* homozygous mice. Those mice were then mated with constitutive *Sos2* (*Sos2 ^−/−^*) null mutant mice(9) to obtain heterozygous Sos1*^fl-Cre/+^* / Sos2*^+/−^* mice. Interbreeding these heterozygous mice, the four genotypes of interest were produced: WT (*Sos1^+-Cre/+-Cre^/ Sos2^+/+^*), SOS1 KO (*Sos1^fl-Cre/fl-Cre^/ Sos2^+/+^)*, SOS2 KO (*Sos1^+-Cre/+-Cre^/ Sos2^−/−^*), and SOS 1/2 DKO (*Sos1^fl-Cre/fl-Cre^/ Sos2^−/−^*)(10).

### Animals and housing conditions

The mice were housed in cages with adequate space and bedding material under specific pathogen-free conditions. They were maintained on a 12-h dark/light cycle at 20-24°C and humidity of 45-65 %. All animal care and experiments adhered to European (2007/526/CE) and Spanish (RD1201/2005 and RD53/2013) regulations and were approved by the Bioethics Committee of the Universidad de Salamanca (#987, #1263). All experiments were performed using male mice only.

### Tamoxifen administration and experimental timeline

TMX (MedChem, #HY-13757A) was administered orally to all experimental groups at a concentration of 40 mg/ml, dissolved in sunflower oil (Sigma, #S50070), for three consecutive days, one dose per day, and PCR monitored *Sos1* loss on days 0, 2, 6, and 11. Mice underwent regular weight and temperature measurements, with the latter recorded using a rectal probe coupled with a Small Animal Physiological Monitoring System (Harvard Apparatus, #75-1500). Finally, mice were euthanized on days 0, 6, and 12.

### Tissue collection and processing

On days 0, 6, and 12, the small intestine, inguinal white adipose tissue (iWAT), and gonadal white adipose tissue (gWAT) were extracted, measured, and weighed for all experimental groups.

### Clinical scoring

The Murine Sepsis Score (MSS) was determined in SOS1/2 DKO mice during TMX treatment based on the criteria described by Sulzbacher *et al*(31).

### Blood collection and biochemical and hematological analyses

Blood samples from all genotypes during TMX treatment were collected from the submandibular vein and subjected to various analyses. Biochemical parameters, including glucose (GLU), Alanine aminotransferase (ALT), Aspartate aminotransferase (AST), and Lactate dehydrogenase (LDH), were measured in serum using the SoptchemTM E7 SP4430 (Arkray). Simultaneously, samples collected in Microvette ethylenediaminetetraacetic acid (EDTA)-coated tubes were analyzed using the HEMAVET 950 (Drew Scientific Inc., Miami Lakes, FL, USA) to assess hematological parameters, such as hemoglobin (Hb), total white blood cells (WBC), red blood cells (RBC), platelets (PLT), and the percentages of neutrophils (NE), lymphocytes (LY), monocytes (MO), eosinophils (EO), and basophils (BA).

### Bacterial culture and identification

Serum samples from all genotypes during TMX treatment were serially diluted and cultured on Luria-Bertani (LB) agar plates at 37 °C for 24 hours. After the incubation, colonies were counted individually for each sample. Subsequently, bacterial load (CFU/mL) was calculated by multiplying the mean colony count by the dilution ratio. Additionally, bacterial load was determined from chopped samples of the heart, lung, liver, and spleen. A biochemical test battery using APIStaph (20500-Biomerieux) was performed to identify bacteria isolated from the serum and organs of SOS1/2 DKO mice.

### Behavioral and metabolic phenotyping of mice in metabolic chambers

Food intake of all experimental groups was measured every two days in individually housed mice. Energy expenditure, respiratory quotient, and locomotor activity were assessed using an indirect calorimetry system (LabMaster, TSE Systems, Bad Homburg, Germany). Rectal temperature was determined using a digital thermometer (VedoBEEP, Artsana, Grandate, Italy). In contrast, skin temperature in the interscapular region surrounding BAT was monitored with an infrared thermal imaging camera (B335 Compact Infrared Thermal Imaging Camera, FLIR, West Malling, Kent, UK). All behavioral and metabolic analyses were conducted at the CIMUS (Centro Singular de Investigación en Medicina Molecular y Enfermedades Crónicas) in Santiago de Compostela.

### Therapeutic interventions

SOS1/2 DKO mice were treated with N-acetylcysteine (NAC, A7250, Sigma-Aldrich) at a concentration of 0,5% (w/v)(32,33) and glucose at 40% (w/v) in drinking water, and a high-fat diet (HFD, ENVIGO MD.06415 (Adjusted Calories Diet (45/Fat) PF1916) was made available ad libitum during TMX treatment. Enrofloxacin (Harmony) was prepared at 50 mg/mL in saline and administered by subcutaneous injection once daily at 3.8 mg/kg body weight. Treatment was carried out for 7 consecutive days, starting on day 5 after TMX administration. During the treatments, SOS1/2 DKO mice underwent weight measurements, temperature monitoring, and blood sample collection for further analysis.

### Glucose and insulin curves

Glucose and insulin tolerance tests (GTT and ITT) were performed following the previously established protocol(34). Mice from all experimental groups treated with TMX for 12 days and without TMX treatment were fasted for 4 hours before the administration of glucose (2g/kg) by oral gavage and for 2 hours before the intraperitoneal insulin (1U/kg, 710014.0, Lilly) injection. Baseline blood glucose levels were assessed before the administration of insulin and glucose. Subsequent measurements were taken at specific time intervals of 15-, 30, 60-, and 120 minutes post-administration. Glucose concentrations were measured using an Accu-Chek Performa blood glucose meter.

### Tissue processing

Paraffin-embedded intestine, iWAT, and gWAT samples were cut and stained with hematoxylin and eosin (H&E) according to the PMC-BEOCyL Unit (Comparative Molecular Pathology-Biobank Network of Oncological Diseases of Castilla y León). Imaging was performed with an Olympus BX51 microscope coupled to an Olympus DP70 digital camera.

### Histological quantifications

The diameter of the adipocytes was measured using stained histological samples acquired at 20x magnification. Ten representative regions per animal were analyzed using the “Adipocyte Tool” macro in ImageJ software (National Institutes of Health, USA).

### Immunohistochemistry and immunofluorescence

Immunohistochemistry stainings of anti-phospho-ERK1/2 (Cell Signaling, #9101, 1:200), anti-phospho-S6 ribosomal protein (Cell Signaling, #5364, 1:1000), anti-Ki-67 (BD Pharmigen, #550609, 1:100) and anti-Cleaved-Caspase-3 (Cell signaling, #9661, 1:400) were performed by the Platform of the IDIBELL Scientific and Technical Services, Comparative Molecular Pathology-Biobank Network of Oncological Diseases of Castilla y León, or in-house as previously described(35).

For immunofluorescence, lysozyme (Lyz) staining (Sigma, #278A-1, 1:200) was performed in-house as previously described(35). Sections were dewaxed, microwaved in citrate buffer (pH 6), and incubated overnight with the primary antibody at 4 °C. Sections were then incubated with the secondary antibody (goat anti-rabbit Alexa 488, Jackson ImmunoResearch, 1:400) and counterstained with nuclear DAPI (Sigma, #D9542,1:1000) for 1 h at room temperature (RT). Finally, sections were mounted with ProLong Diamond anti-fading reagent (Life Technologies, P36970) and images were acquired using the Thunder Imager Tissue (Leica DM6B, Wetzlar, Germany) with a 20x objective.

### Staining quantifications

The quantifications of phospho-ERK1/2, phospho-S6, and Ki-67 in the small intestine were performed using R (version 4.3.1; R Core Team, 2023) in RStudio (version 2022.02.1+461; RStudio Team, 2022). The R-script detects the immunohistochemistry signal (brown pixels) and calculates the signal by dividing brown pixels/ blue pixels representing the nucleus to ensure accuracy. For each animal, 10 representative regions from histology samples acquired at x20 magnification were selected for quantification.

The image J 1.53c software was utilized to quantify lysozyme staining in the small intestine. This process involved identifying pixel areas corresponding to these targets in six images per animal, acquired with the fluorescence microscope (Leica DM6B, Wetzlar, Germany). Each image contained 10 regions at 20x magnification.

The quantification of cleaved caspase-3 staining in the small intestine was performed using QuPath-0.5.1 software(36). QuPath was used to detect DAB-positive and DAB-negative cells. The percentage of positive cells relative to total cells was then calculated using R. Quantification was based on pictures obtained from all liver tumors at 20x magnification.

### Intestine stem cell extraction and quantification

Small intestines from all experimental groups were carefully cleaned and opened longitudinally to isolate intestinal stem cells following established protocols(37,38). The tissue fragments were pipetted up and down in ice-cold PBS until the solution turned clear. Once the solution cleared, the pieces were incubated in 2.5 mM EDTA in PBS for 30 min at 4 °C. After incubation, the supernatant was removed, and the small pieces were rinsed in ice-cold PBS containing 10% FBS. Crypts were released by gentle shaking, and this step was repeated several times to increase yield. The suspension was filtered through a 70 μm strainer (Corning, #CLS352350) and centrifuged at 1500 rpm for 2 min at 4 °C. Crypts were then incubated in DMEM/F12 for 30 min at 4 °C, washed with ice-cold PBS, and centrifuged at 1500 rpm for 3 min at 4 °C. Crypts were then resuspended in TrypLE supplemented with 10 μM Y-27632 (Medchem, #HY-10583) and 1 µg/mL DNase, and incubated for 5 min at 37 °C. The reaction was quenched by adding 200 μL of FBS, followed by dilution in 30 mL of ice-cold PBS, filtration through a 70 μm strainer, and centrifugation at 1500 rpm for 3 min. The resulting IEC pellet was washed in PBS and stained with appropriate fluorescently labeled antibodies. Stained IECs were analyzed on a BD FACSAria III flow cytometer, and stem cell populations were quantified based on DyLight^TM^ 488-Lgr5 (Origene, #TA400002) and APC-CD44 (BD, #559250) expression.

### Isolation of peritoneal immune cells

Peritoneal immune cells were isolated using a previously established peritoneal lavage protocol(39). Briefly, mice were euthanized, sprayed with 70% ethanol, and mounted on a Styrofoam block. The peritoneum was exposed by cutting the outer skin and gently retracting it. 5 mL of ice-cold PBS supplemented with 3% FBS was injected into the peritoneal cavity using a 25G needle, followed by a gentle massage to dislodge cells. The lavage fluid was collected, taking care to minimize blood contamination. The collected cell suspension was centrifuged at 300 x g for 5 min at 4 °C, the supernatant was discarded, and the pellet was resuspended in PBS for staining. Peritoneal cavity cells were stained with anti-mouse Gr1-FITC (Biolegend, #108406), CD45R/B220-PerCP Cy5.5 (BD, #561101), CD3e-APC (Invitrogen, #17-0031-82), and analyzed by flow cytometry (BD FACSAria III).

### Establishment of intestinal organoids

Organoids were established following a previously described protocol(37). Freshly isolated small intestines from WT, SOS1*^fl/fl^*/ SOS2 KO, and C57BL/6J x BALB/c-Tg(CMV-cre)1Cgn/J(40) mice were longitudinally opened and washed three times with ice-cold PBS. The intestines were then chopped into pieces of 2 to 5 mm and placed in ice-cold PBS. The chopped pieces were pipetted up and down in ice-cold PBS until the solution turned clear. Once the solution cleared, the pieces were incubated in 2.5 mM EDTA in PBS for 30 minutes at 4°C. After incubation, the supernatant was removed, and the small pieces were rinsed in ice-cold PBS containing 10% FBS (Gibco, #A5256701). The intestinal pieces were gently shaken to release free crypts. This step was repeated several times to obtain a higher number of crypts. The suspension was then filtered through a 100 µm mesh. Finally, a total of 200 isolated crypts were counted and embedded in 20 µL of undiluted Matrigel (Cultek, #356237). The culture medium used for the crypts consisted of advanced DMEM/F12 media (#12634-010, Gibco) supplemented with Pen/Strep (Gibco, #15140122), HEPES (Gibco, #15630-080, 1X), NAC (Sigma-Aldrich, #A7250, 1mM), B27 (Gibco, #17504-044, 1X), EGF (Sigma, #E9644, 50ng/mL), Nogging (MedChem, #HY-P7086, 100ng/mL), R-spondin (MedChem, #HY-P72784A, 500ng/mL), and Y-27632 inhibitor (MedChem, #HY-10583, 10 µmol/L).

### Generation of L-WRN conditioned medium protocol

L-WRN conditioned medium was generated from L-WRN cells (ATCC CRL-3276) as previously described(41). These cells were cultured in Advanced DMEM/F12 medium supplemented with 10% FBS, 1% GlutaMAX, and 1% P/S under standard conditions (37 °C, 5% CO₂). Once cells reached approximately 70-80% confluency, the culture medium was replaced with serum-free Advanced DMEM/F12, and cells were incubated for an additional 7 days before collecting the conditioned medium. The collected supernatant was then centrifuged at 300 × g for 5 min to remove cell debris, filtered through a 0.22 µm PES filter, and stored at 4 °C for up to one week or at −80 °C for long-term storage.

### Generation of organoid-derived intestinal epithelial monolayers

Monolayers of intestinal epithelial cells were formed from organoids using a method established in earlier studies(42). Organoid cultures were enriched for stem cells through 4-5 passages using the organoid culture medium described above. Mature organoids were then mechanically dissociated using a 30G syringe. This procedure yielded a suspension of single cells or small cell clusters. Cells were seeded onto 5% Matrigel-coated cell culture inserts (Corning, #356237) at a density of 10^⁵^ cells in an initial volume of 10-20 µL. Cells were allowed to adhere for 1 h before the medium was added to a final volume of 100 µL. The upper chamber medium did not contain L-WRN-conditioned medium, while the lower chamber (basal compartment) received medium supplemented with L-WRN to generate a chemotactic gradient.

### FITC-DEAE-Dextran permeability assays

#### *In vivo* intestinal permeability

Intestinal permeability was assessed as previously described(43). Mice of all genotypes undergoing TMX treatment received orally 150 µL of FITC-DEAE-Dextran (4 kDa; 80 mg/mL). Blood samples were collected immediately before and 4 h after gavage using serum collection tubes. Plasma was isolated, diluted in PBS, and fluorescence was measured in black 96-well plates using a microplate reader (Infinite® 200 PRO, Tecan) at 495/520 nm (excitation/emission). Measurements were performed in duplicate or triplicate.

#### *In vitro* permeability assay

Epithelial permeability *in vitro* was evaluated using IEC cultured on cell culture inserts after 6 days of treatment with 0.4 µM hydroxytamoxifen (4-OHT, Sigma-Aldrich, #H6278) as previously described(42). Briefly, apical and basolateral compartments were washed with phenol red-free DMEM supplemented with 1% HEPES. FITC-DEAE-Dextran (0.5 mg/mL, 200 µL) was added to the apical compartment, and phenol red-free DMEM (600 µL) to the basolateral compartment. Cells were maintained at 37 °C, and basolateral samples were collected at defined time points up to 2 h. Fluorescence was measured at 495/520 nm using a microplate reader. FITC-Dextran fluxed across inserts with or without Matrigel coating was used as a control.

#### Surgical technique and orthotopic interventions

SOS1/2 DKO mice on day 6 of TMX treatment were weighed and anesthetized with inhaled isoflurane (Vetflurane, #03597133058451), followed by a preoperative subcutaneous injection of 0,1 mg/kg of buprenorphine (Bupaq, VetViva). Mice were positioned on a warming pad and covered with sterile drapes. A midline laparotomy was performed to expose the small intestine. Treatments were then administered directly into the intestinal lumen using a 30G syringe, with a final volume of 300 µL for all conditions: (**1**) Intestinal organoids suspended in 5% Matrigel (Corning, #356237) in PBS(44); (**2**) 5% Matrigel (Corning, #356237) in PBS; (**3**) 30% L-WRN-conditioned medium in DMEM/F12. Following treatment administration, the abdominal musculature and skin were closed using Vicryl sutures (Mersilk, #W528). Mice were monitored during recovery from anesthesia and maintained on a soft gel diet for several days post-surgery.

#### qRT-PCR

Total RNA was extracted from the IECs using NZYol reagent (nzytech, #MB18501) and analyzed by quantitative real-time polymerase chain reaction (qRT-PCR) using the Luna Universal One-Step RT-qPCR Kit (BioLabs, #E3005L). According to the manufacturer’s instructions, the qRT-PCR reactions were performed on a Quant Studio 5 Pro Real-Time PCR system (Applied Biosystems, #A43172). Relative mRNA expression levels were calculated using the 2-ΔΔCt method after normalization to the housekeeping gene β2-microglobulin. Primers used are listed in Supplementary Table 1.

#### Statistical analysis

The experiments were conducted with a minimum of three independent biological replicates for each condition. The specific number of experimental replicates (n values) for each condition is provided in the respective figure legends. Statistical analyses were performed using GraphPad Prism 8. To determine statistical significance, data were analyzed using either one-way or two-way ANOVA, as appropriate. One-way ANOVA was used for comparisons between groups at a single time point or without a time component, followed by a post hoc test for multiple comparisons. Two-way ANOVA was employed to assess the effects of both genotype and treatment over time. To account for multiple comparisons, either the Bonferroni test or the Sidak method was applied. All data are presented as the mean value along with the standard error of the mean (s.e.m.). Differences between groups were considered statistically significant when the p-value was less than 0.05 (p<0.05).

Insulin and glucose responses were further analyzed by calculating the area under the curve (AUC) for each curve using GraphPad Prism, providing an integrated measure of overall response over time.

Survival analyses were performed using Kaplan-Meier survival curves, with differences between groups evaluated by the log-rank (Mantel-Cox) test. Hazard ratios (HR) with 95% confidence intervals were calculated to estimate the relative risk between experimental conditions.

## RESULTS

### Loss of SOS1 and SOS2 leads to dramatic weight loss, reduction of fat tissues, and sudden death

Following three consecutive daily oral gavages of TMX (40 mg/mL per dose) in male mice, SOS1/2 DKO mice exhibited continuous weight loss, reaching 30% of their total body weight, and 100% lethality within 12 days. In contrast, single-KO counterparts and WT littermates showed no change (Figs. 1A and B).

**Figure 1.**
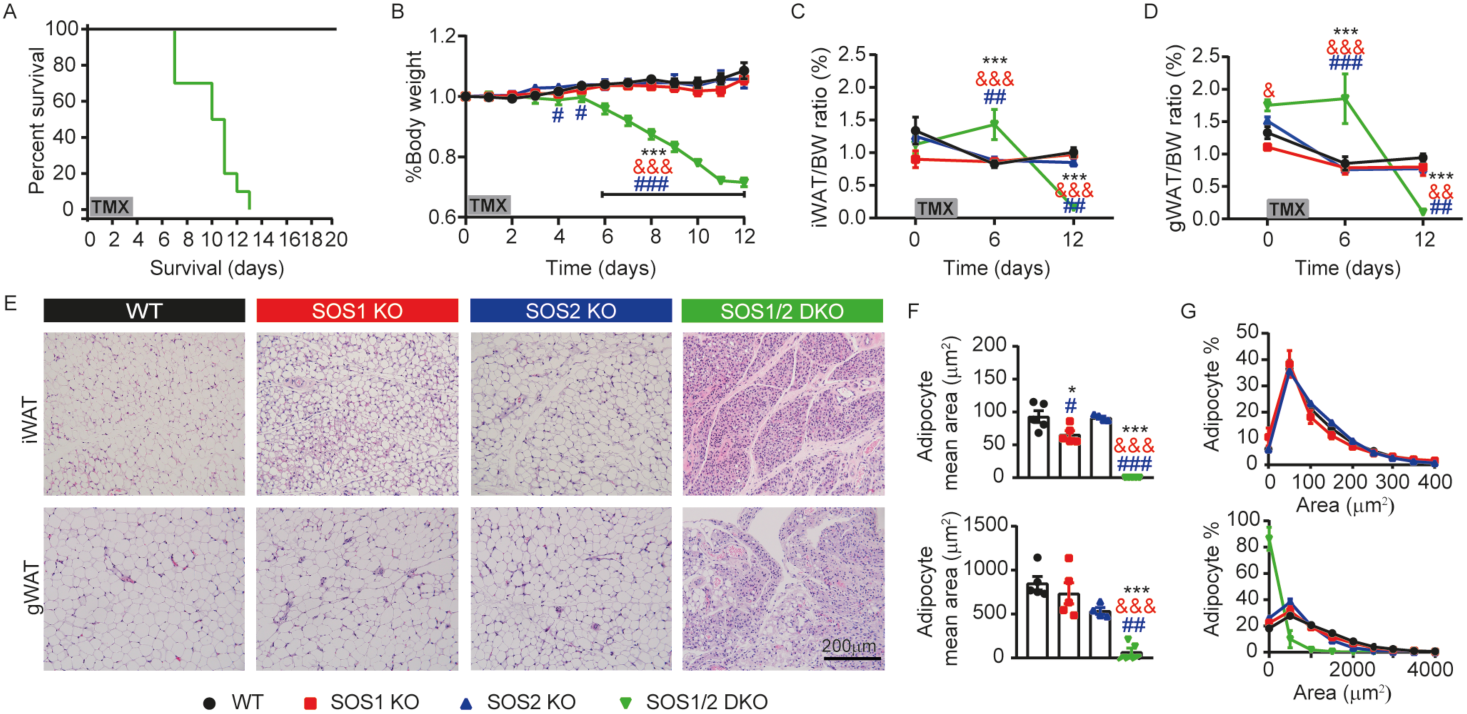
Combined genetic ablation of SOS1 and SOS2 induces severe lipoatrophy and lethality. (A) Survival curves of WT, SOS1 KO, SOS2 KO, and SOS1/2 DKO mice after TMX treatment. (B) Evolution of the body weight in mice of all genotypes after TMX treatment. Body weight values were normalized on the first day before TMX administration (day 0). (C-D) Changes in adipose tissue after TMX administration. The weight of iWAT (C) and gWAT (D) was normalized against body weight (BW) on days 0, 6, and 12. (E) Representative H&E-stained sections of iWAT (upper sections) and gWAT (lower sections) from WT, SOS1 KO, SOS2 KO, and SOS1/2 DKO mice on day 12 of TMX treatment. (F-G) Adipocyte size analysis 12 days after TMX treatment across genotypes. (F) Median adipocyte area in iWAT (upper panel) and gWAT (lower panel). (G) Percentage distribution of adipocytes according to cell area in iWAT (upper panel) and gWAT (lower panel). Data expressed as the mean ± s.e.m. for at least n=5 for each genotype per day. Statistical analysis was performed using one-way or two-way ANOVA, as appropriate, with Tukey’s post hoc test. Significance indicators: * vs. WT, & vs. SOS1 KO and # vs. SOS2 KO; *, &, #, p<0.05; &&, ##, p<0.01; ***, &&&, ### p<0.001.

The significant weight loss prompted us to examine the status of adipose depots. Notably, mice lacking both SOS proteins showed a marked reduction in iWAT and gWAT weight, consistent with their overall weight loss. In contrast, WT, SOS1 KO, and SOS2 KO mice displayed no significant changes in fat pad content between days 0 and 12 (Figs. 1C and 1D).

To determine whether SOS1 and SOS2 influence WAT remodelling, we analysed the histological structure of iWAT and gWAT. Interestingly, deletion of SOS1 at room temperature (without cold exposure) resulted in pronounced alterations in adipose tissue architecture, shifting it towards a more thermogenic profile. Specifically, H&E staining at day 12 revealed an increase in multilocular adipocytes and a reduction in adipocyte size in SOS1 KO mice. In contrast, adipose tissue from SOS1/2 DKO mice at day 12 showed severe adipocyte atrophy with reduced unilocular vacuoles by H&E (Figs. 1E-G).

### SOS1/2 DKO mice exhibit systemic metabolic failure and liver dysfunction

To better understand the weight loss observed in SOS1/2 DKO mice and the histological changes in the iWAT of SOS1 KO mice on day 12, we investigated their energy metabolism. WT, SOS1 KO, SOS2 KO, and SOS1/2 DKO mice were housed in metabolic chambers during TMX treatment, and their metabolic rates were assessed via indirect calorimetry. Surprisingly, no significant differences were observed in daily food intake or overall energy expenditure among the genotypes (Figs. 2A and B). However, SOS1/2 DKO mice exhibited reduced locomotor activity and lower body temperature (Figs. 2C and D). Additionally, SOS1/2 DKO mice showed a lower respiratory quotient (RQ: VCO2/VO2), indicating a metabolic shift toward the utilization of lipids rather than carbohydrates to meet energy demands (Fig. 2E).

**Figure 2.**
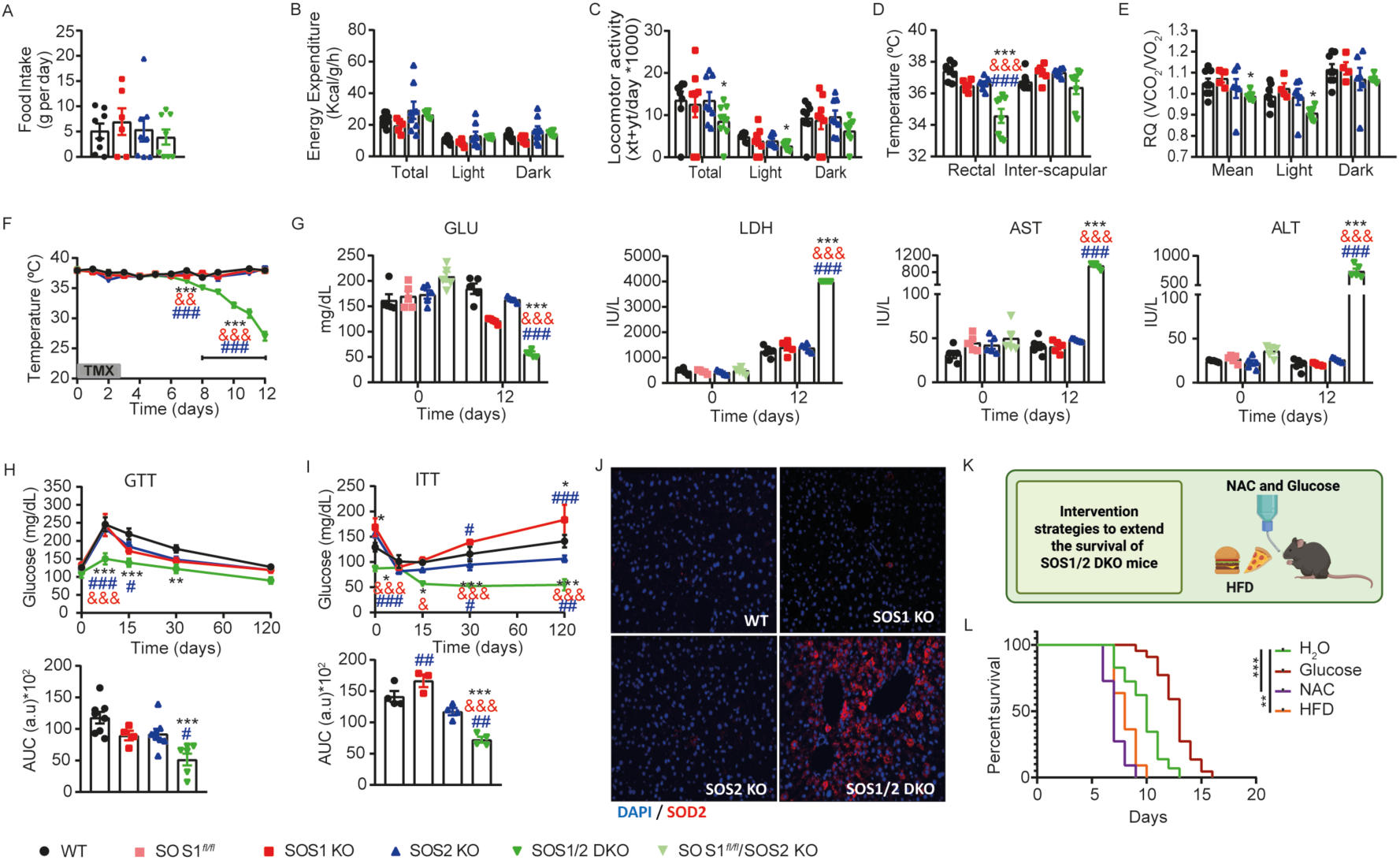
Simultaneous genetic ablation of SOS1 and SOS2 causes hypoglycemia and hepatic dysfunction. (A-D) Metabolic cage analysis of WT, SOS1 KO, SOS2 KO, and SOS1/2 DKO mice 12 days after TMX treatment. Bar graphs show food intake (A), energy expenditure. (B), and locomotor activity (C) during the total, light, and dark phases, body temperature (D; rectal and interscapular), and respiratory quotient (E) during the mean, light, and dark phases. (E) VCO2 represents the volume of CO2, and VO2 represents the volume of oxygen. (F) Evolution of body temperature of WT, SOS1 KO, SOS2 KO, and SOS1/2 DKO mice during treatment. (G) Glucose (GLU), Lactate dehydrogenase (LDH), Aspartate aminotransferase (AST), and Alanine aminotransferase (ALT) were measured in the serum of all experimental groups on days 0 and 12. (H) A glucose tolerance test (GTT) was performed 12 days after TMX treatment in the indicated genotypes. The upper panel shows blood glucose levels over time, and the lower panel shows quantification of the area under the curve (AUC). (I) An insulin tolerance test (ITT) was performed 12 days after TMX treatment in the indicated genotypes. The upper panel shows blood glucose levels over time, and the lower panel shows quantification of the area under the curve (AUC). (J) SOD2 immunostaining was performed on all genotypes on day 12 of TMX treatment. (K) Schematic representation of the intervention strategies to extend the survival of SOS1/2 DKO mice, glucose, and N-acetylcysteine (NAC) in drinking water and high-fat diet (HFD). (L) Survival rates for all treatment conditions in SOS1/2 DKO mice are represented in the Kaplan-Meier graph. Data expressed as the mean ± s.e.m. for at least n=4 for each genotype per day. Statistical analysis was performed using one-way ANOVA, with Tukey’s post hoc test. Significance indicators: * vs. WT, & vs. SOS1 KO and # vs. SOS2 KO; *, &, #, p<0.05; **, &&, ##, p<0.01; ***, &&&, ### p<0.001.

Continuous monitoring of body temperature throughout TMX treatment revealed a marked, progressive decline in SOS1/2 DKO mice that closely paralleled their severe weight loss. In contrast, WT, SOS1 KO, and SOS2 KO mice maintained stable body temperatures over the entire treatment period (Fig. 2F). Alongside the development of hypothermia, SOS1/2 DKO mice also exhibited significantly reduced blood glucose levels following TMX administration (Fig. 2G and Suppl. Fig. S1C). To further characterize glucose homeostasis, a GTT was performed before (Suppl. Fig. S1A) and after 12 days of TMX treatment. Post-treatment GTT revealed markedly lower blood glucose levels in SOS1/2 DKO mice at 15, 30, and 60 minutes following glucose administration compared with the other genotypes (Fig. 2H). A similar pattern was observed in the ITT, with SOS1/2 DKO mice exhibiting significantly lower glycemia at 15, 30, and 60 minutes after insulin injection (Fig. 2I), indicating an overall state of hypoglycemia and altered glucose handling.

In addition to these systemic metabolic defects, biochemical analyses revealed a significant increase in circulating markers of liver injury, including AST, ALT, and LDH, in SOS1/2 DKO mice (Fig. 2G). Given the elevation of hepatic dysfunction markers, we next examined oxidative stress in the liver by immunostaining for superoxide dismutase 2 (SOD2) in liver sections from SOS1/2 DKO mice at day 12. This analysis revealed increased SOD2 expression in SOS1/2 DKO livers compared with controls (Fig. 2J), suggesting enhanced oxidative stress and liver damage associated with the combined loss of SOS1 and SOS2.

To determine if these metabolic alterations were the primary cause of death, we tested several intervention strategies to extend the survival of SOS1/2 DKO mice, including glucose supplementation, administration of an HFD, and the antioxidant compound NAC (Fig. 2K). Despite the severe hypoglycemia observed in SOS1/2 DKO mice, oral glucose supplementation modestly increased the survival of SOS1/2 DKO mice, but it did not stop SOS1/2 DKO mice from dying. To revert the extensive fat loss in SOS1/2 DKO mice, we administered an HFD; however, this treatment not only failed to improve survival but also led to earlier death. Similarly, NAC, which we hypothesized could mitigate oxidative stress and prolong the lifespan, failed to prevent mortality and instead accelerated death (Fig. 2L).

Given the ineffectiveness of the aforementioned metabolic interventions and the rapid deterioration of the animals, we hypothesized that the observed metabolic collapse (lipoatrophy and hypoglycemia) might instead be a systemic consequence of failure at the level of specific organ(s).

### Genetic disruption of both SOS GEFs leads to impaired intestinal homeostasis

Previous studies have highlighted the critical role of the ERK/MAPK pathway in regulating cell proliferation in the small intestine(27,28). Based on this, ablation of both SOS GEFs could be expected to impair intestinal epithelial growth. As none of the previous analyses or interventions fully explained or prevented the mortality observed in SOS1/2 DKO mice, we next examined their intestinal structure and cellular proliferative capacity.

Before TMX administration, no overt differences were observed between genotypes regarding intestinal morphology, cellular composition, or signaling pathway activity, suggesting baseline equivalence (Suppl. Fig. S2). Early after TMX induction, by day 5, proliferation and ERK phosphorylation were reduced in SOS2 KO and SOS1/2 DKO mice, and a decrease in ISC numbers was observed in SOS1/2 DKO intestines (Suppl. Fig. S3). These early molecular and cellular changes preceded the more pronounced, overt macroscopic defects that were clearly visible on day 12 of TMX treatment, when the small intestine of SOS1/2 DKO mice exhibited luminal distension, a trend toward length shortening, and loss of tissue integrity (Fig. 3A, black arrows). Quantitative analysis confirmed that both SOS1/2 DKO and SOS1 KO mice displayed intestinal shortening, with the effect being most severe in SOS1/2 DKO animals; SOS1 KO intestines were also significantly shorter than those of SOS2 KO mice (Fig. 3A).

**Figure 3:**
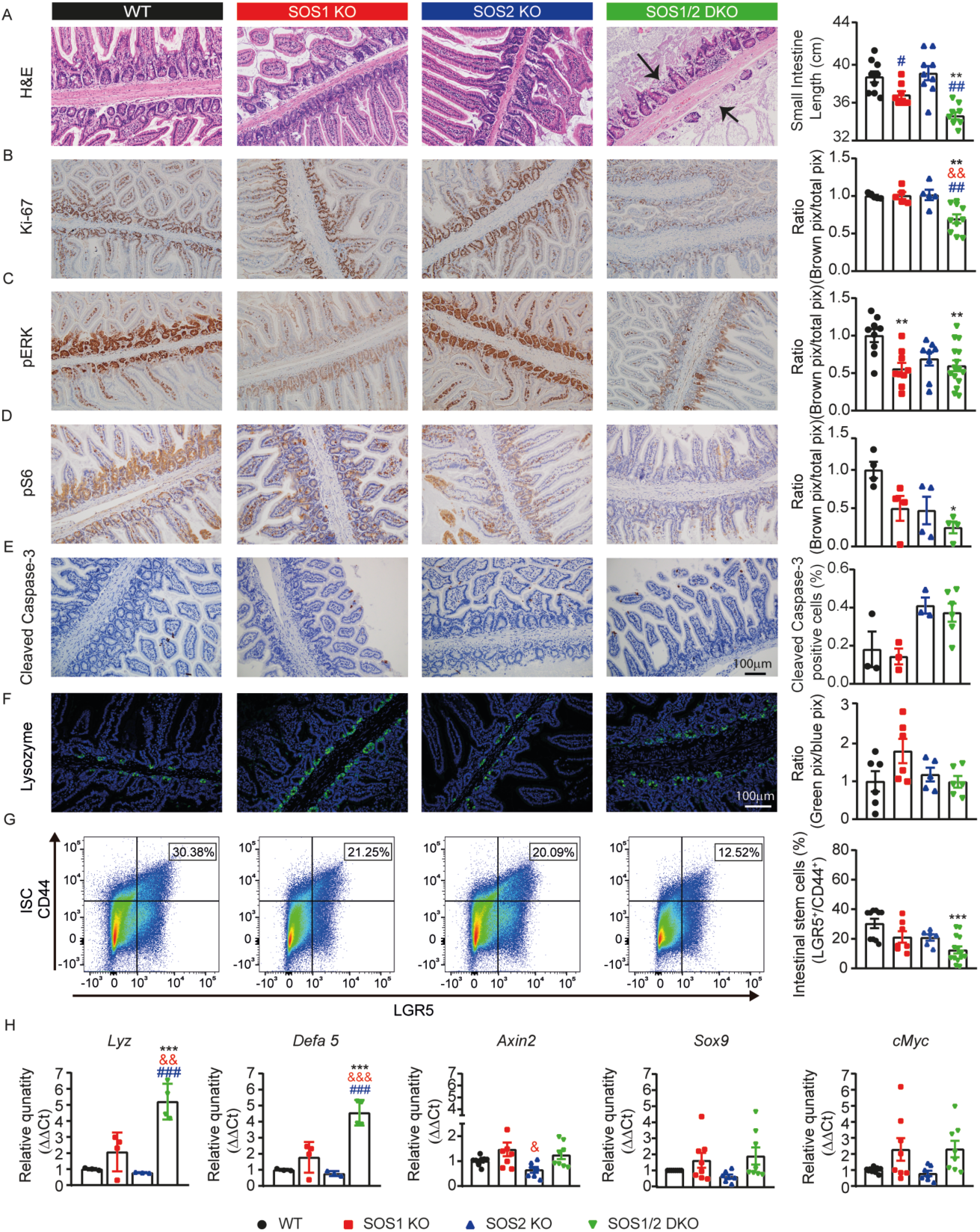
SOS1 and SOS2 genetic disruption results in reduced proliferation, decreased ERK and S6 phosphorylation, loss of intestinal structure, and decreased ISC numbers. (A) Representative hematoxylin and eosin (H&E)-stained sections of the small intestine from all experimental groups on day 12 after TMX treatment, and Small Intestine Length quantification from each experimental group on day 12 after TMX treatment. (B-E) Representative immunohistochemistry images showing Ki-67 (B), pERK (C), pS6 (D), and cleaved caspase-3 (E) staining in WT, SOS1 KO, SOS2 KO, and SOS1/2 DKO mice 12 days after TMX treatment, with quantification. (F) Representative lysozyme immunofluorescence images of the small intestine of the four genotypes under study on day 12 after TMX treatment, with quantification. (G) Representative FACS profiles of LGR5^+^/CD44^+^ expression and the frequency of LGR5^+^/CD44^+^ Intestinal Stem Cells (ISC) quantification from each experimental group on day 12 after TMX treatment. (H) mRNA expression levels of the indicated genes in stem cells isolated from the small intestine of WT, SOS1 KO, SOS2 KO, and SOS1/2 DKO mice on day 12. Data expressed as the mean ± s.e.m. for at least n=3 for each genotype. Statistical analysis was performed using one-way ANOVA, with Tukey’s post hoc test. Significance indicators: * vs. WT, & vs. SOS1 KO and # vs. SOS2 KO; &, # p<0.05; **, &&, ## p<0.01; ***, &&&, ### p<0.001.

Given these structural alterations, we next assessed pathways associated with epithelial proliferation. SOS1/2 DKO mice exhibited a reduced number of Ki-67^+^ proliferating cells on day 12 compared with the other genotypes. In parallel, ERK and S6 phosphorylation were decreased relative to WT mice (Figs. 3B-D). To determine whether the loss of intestinal tissue was driven by active cell death, we assessed apoptosis via cleaved caspase-3 staining. We detected no significant increase in apoptotic markers in SOS1/2 DKO crypts compared to controls (Fig. 3E). This absence of massive cell death, combined with the proliferation block, suggests that the intestinal atrophy is primarily driven by a failure of epithelial renewal and crypt exhaustion, rather than by acute cytotoxicity. In contrast, SOS1 KO mice maintained stable numbers of Ki-67^+^ cells over time, potentially compensating for the reduced ERK phosphorylation observed on day 12 (Figs. 3B-D).

Maintaining intestinal homeostasis relies not only on proliferation but also on the harmonized presence of diverse cell populations, including Paneth cells and ISC. Consistent with this, SOS1 KO mice showed an increase in Paneth cells as indicated by lysozyme staining. In contrast, SOS1/2 DKO mice did not display this increase in Paneth cells (Fig. 3F). Together with the reduced proliferation observed earlier, they exhibited a diminished number of ISCs on day 12, indicating impaired regenerative capacity (Fig. 3G).

Moreover, the expression of Paneth cell-associated genes (*Lyz* and Defensin Alpha 5 (*Defa5*)) showed an upward trend in SOS1 KO and SOS1/2 DKO ISCs on day 12 compared to WT and SOS2 KO. Conversely, genes linked to the Wnt pathway (*Axin2*, *Sox9*, and *cMyc*), key regulators of intestinal regeneration, were not significantly overexpressed in SOS1 KO and SOS1/2 DKO mice, although they showed a tendency to rise (Fig. 3H).

### Gut permeability and bacterial translocation in SOS1/2 DKO mice

Given the observed intestinal damage, we opted to employ a 4-kDa FITC-DEAE-Dextran (FITC-Dextran) assay to assess gut permeability. Upon oral administration, the FITC-Dextran fluorescent marker traverses the gastrointestinal tract and readily permeates passively through the intestinal epithelium (Fig. 4A). The quantification of FITC-Dextran concentration in plasma, facilitated by a fluorimeter, serves as a reliable metric for assessing the paracellular permeability of the intestinal epithelium(43).

**Figure 4:**
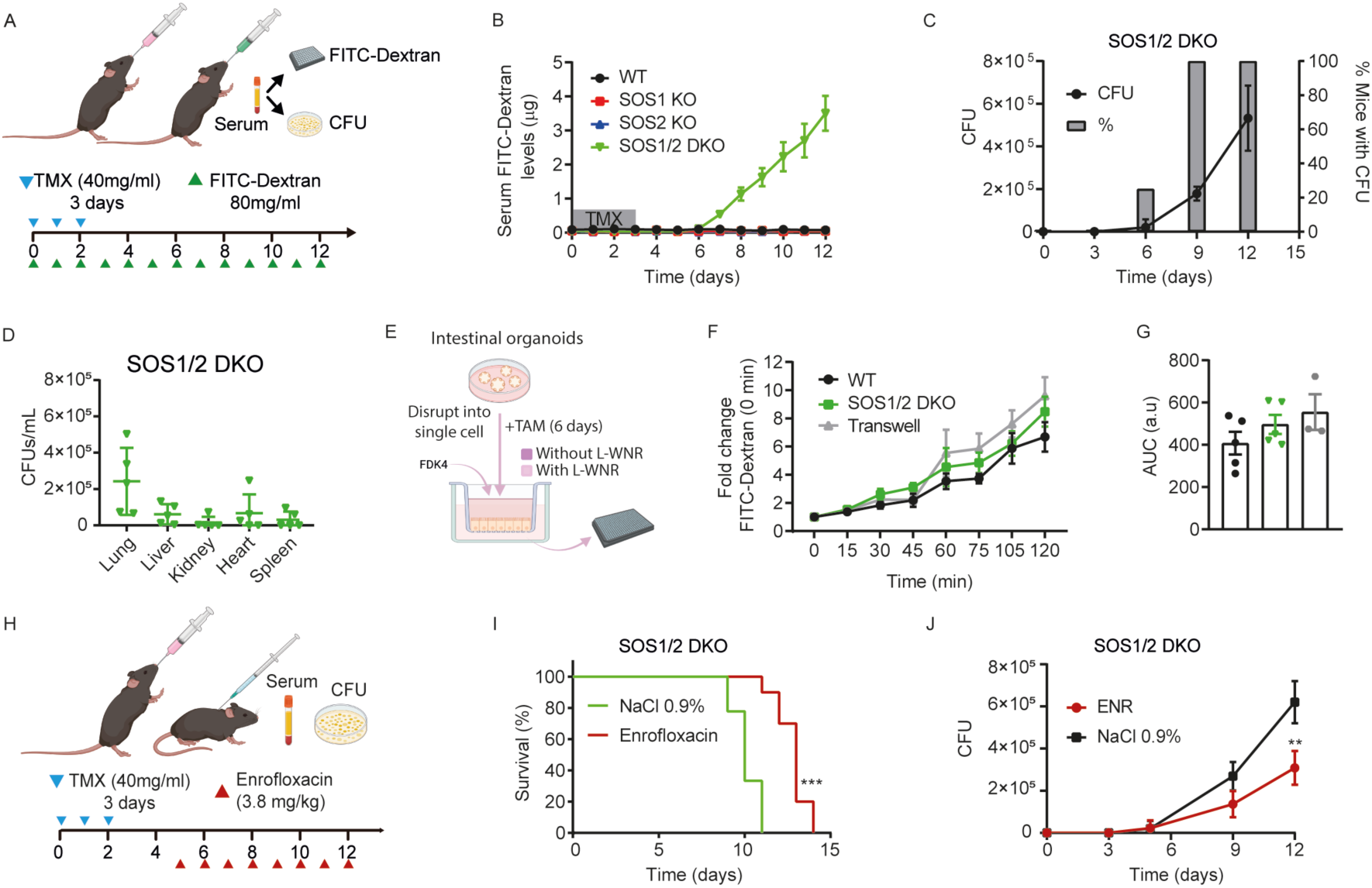
Combined loss of SOS1 and SOS2 significantly increases gut permeability and bacterial translocation. (A) Schematic representation of the experimental setup to assess intestinal permeability and bacterial translocation. TMX (40 mg/mL) was administered orally for 3 consecutive days to induce gene deletion, followed by daily oral gavage with FITC-DEAE-Dextran (FITC-Dextran, 4 kDa, 80 mg/mL). (B) Serum and blood samples were collected for analysis. Serum levels of FITC-Dextran were measured by fluorometry to evaluate intestinal barrier integrity. (C) Quantification of bacterial colony-forming units (CFUs) in blood from SOS1/2 DKO mice over time. The line represents the number of CFU over the TMX treatment, and the bars indicate the percentage of mice with detectable bacteremia. (D) Quantification of CFUs in organs from SOS1/2 DKO mice on day 12 post-TMX treatment. (E) Schematic representation of the experimental setup to assess intestinal permeability *in vitro* using intestinal organoids. *Created with BioRender.com.* (F-G) Media was collected every 15 min, and FITC-Dextran levels were measured by fluorometry to evaluate intestinal barrier integrity (F) together with area under the curve quantification (G). (H) Schematic representation of the experimental setup of the antibiotic treatment experiment. SOS1/2 DKO mice received either NaCl 0.9% or enrofloxacin (3.8 mg/kg, subcutaneously) daily, starting after TMX induction. (I) Kaplan-Meier survival curves of SOS1/2 DKO mice treated with enrofloxacin (ENR) or vehicle (0.9% NaCl). (J) Quantification of CFUs in blood from SOS1/2 DKO mice treated with ENR or NaCl over time. Data expressed as the mean ± s.e.m. for at least n=3 for each genotype per day. Statistical analysis was performed using two-way ANOVA, with Tukey’s post hoc test and comparison of survival curves with the Log-rank (Mantel-Cox) test. Significance indicators: * vs. untreated SOS1/2 DKO mice. **, p<0.01; ***, p<0.001.

SOS1/2 DKO mice exhibited elevated serum FITC-Dextran levels from day 6 to day 12 of TMX treatment, indicating increased gut permeability (Fig. 4B). This finding correlates with histological evidence of intestinal damage (H&E staining), supporting a direct link between structural disruption and functional leakiness. Furthermore, bacterial presence was detected in the serum of peripheral blood, which started to appear on day 6 of TMX treatment and kept augmenting over time (Fig. 4C). Consistent with a septic phenotype, we detected a massive bacterial burden in the liver, lung, heart, and spleen of SOS1/2 DKO mice on day 12 after treatment (Fig. 4D), demonstrating active bacterial translocation and systemic dissemination.

An APIStaph (20500-Biomerieux) test revealed that the bacteria found in SOS1/2 DKO mice were *Staphylococcus lentus*. A Gram-positive, coagulase-negative bacterium, resistant to novobiocin, with positive oxidase activity, belonging to the genus *Staphylococcus*, which consists of clustered *cocci. Staphylococcus lentus*(45). It is part of the microbiota in the intestine and skin of mice, where it can experience a significant increase in some cases(46).

To further investigate epithelial integrity at the cellular level, intestinal organoids derived from WT and SOS1/2 DKO mice were cultured up to 4 passages and then disrupted and seeded onto the cell culture inserts to assess monolayer formation and barrier function (Fig. 4E). To better mimic the gradients of biochemical factors generated *in vivo* by the stem cell niche, the basolateral compartment was filled in with a medium containing EGF, Noggin and R-Spondin, while the apical compartment was filled in with basic-medium (not supplemented with previously mentioned factors)(42). Following 6 days of 4-OHT treatment, SOS1/2 DKO cells showed a non-significant trend toward increased permeability to the FITC-Dextran fluorescent marker across the cell culture insert, which may suggest impaired monolayer formation compared to WT controls. This *in vitro* result supports the *in vivo* observations of disrupted epithelial barrier integrity in SOS1/2 DKO mice (Figs. 4F and G).

To mitigate bacterial burden, we administered an antibiotic (enrofloxacin) to the SOS1/2 DKO mice on day 5 after TMX induction and continued daily for 7 days (Fig. 4H). Although this regimen extended lifespan and reduced bacterial load (Figs. 4I and J), it did not improve gut permeability and failed to prevent mortality.

### Disruption of SOS1 and SOS2 induces severe septicemia in mice

To evaluate the consequences of increased gut permeability and bacteremia, we assessed the sepsis score in SOS1/2 DKO mice. Sepsis is a severe medical condition marked by an uncontrolled and dysregulated systemic response to infection that can result in organ dysfunction and, in severe cases, sudden death (47), which aligns with the mortality seen in our SOS1/2 DKO mice. SOS1/2 DKO mice showed a progressive rise in sepsis score starting on day 5 after TMX treatment (Fig. 5A). In parallel, serum markers of liver injury (serum ALT and AST) increased concurrently with the Murine Sepsis Score, showing a strong positive correlation (Fig. 5B and C).

**Figure 5:**
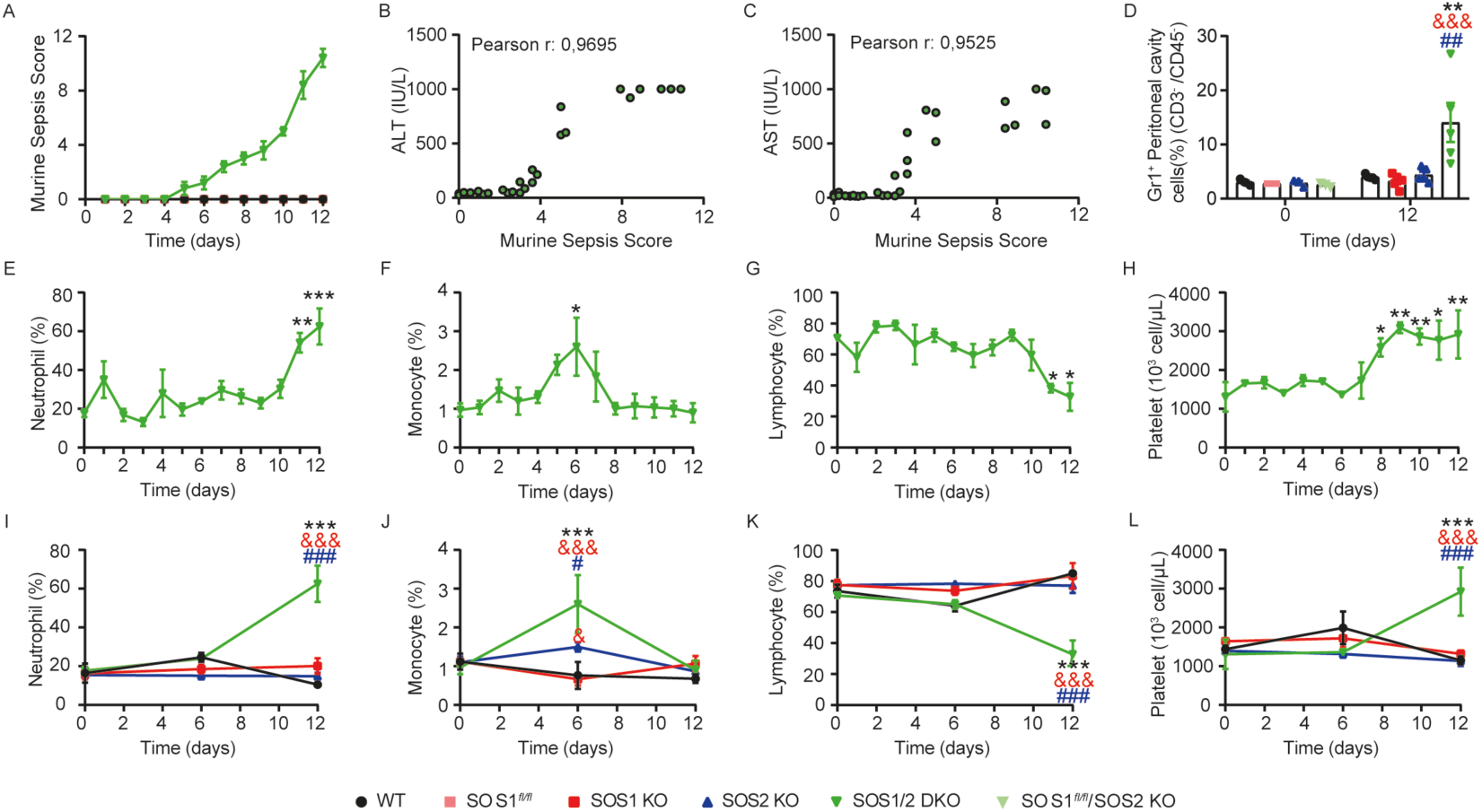
SOS1/2 DKO mice exhibit marked septicemia, liver dysfunction, and enhanced neutrophil infiltration. (A) The Murine Sepsis Score (Methods) was evaluated daily following TMX administration. (B-C) The levels of Serum Alanine aminotransferase (ALT) (B) and Aspartate aminotransferase (AST) (C) levels were measured in SOS1/2 DKO mice and plotted against the Murine Sepsis Score. Pearson correlation coefficients are indicated (Pearson r). (D) The percentage of Gr1^+^ cells (CD3^−^ and CD45^−^) in peritoneal lavage was assessed on days 0 and 12 of TMX treatment in all genotypes. (E-L) Immune profiling of SOS1/2 DKO mice over time (E-H) and across genotypes (I-L) at days 0, 6, and 12 during TMX treatment includes: neutrophils, monocytes, lymphocytes, and platelets. Data expressed as the mean ± s.e.m. for at least n=3 for each genotype per day. Statistical analysis was performed using one-way ANOVA, with Tukey’s post hoc test. Significance indicators for panels E-H: *, p<0.05; **, p<0.01; ***, p<0.001 indicate comparisons with day 0 of SOS1/2 DKO mice. Significance indicators for panels D and from I to L: * vs. WT, & vs. SOS1 KO, and # vs. SOS2 KO; &, # p<0.05; ## p<0.01; ***, &&&, ### p<0.001.

Analysis of peritoneal lavage fluid revealed a marked accumulation of Gr1^⁺^ cells in SOS1/2 DKO mice compared with WT and single KOs, indicating enhanced immature myeloid cells infiltration at the site of infection (Fig. 5D) and a reduction of CD11b^+^ cells (Suppl. Fig. S4). Moreover, hematological profiling further demonstrated a biphasic immune response: an early pro-inflammatory phase marked by transient monocytosis (day 6), followed by a robust neutrophil expansion (Figs. 5E and F). This was subsequently replaced by an immunosuppressive phase, characterized by a sharp reduction in lymphocytes and the development of thrombocytosis with progressively rising platelet counts (Figs. 5G-H). Importantly, these hematological alterations were specific to SOS1/2 DKO mice and were absent in WT or single KO cohorts (Figs. 5I-J and Suppl. Table S2).

### Investigating the therapeutic potential of WT intestinal organoids in SOS1/2 DKO mice

As conventional treatments proved ineffective, we pursued a regenerative strategy inspired by previous reports of successful intestinal engraftment(44). In addition to implanting WT intestinal organoids to evaluate their capacity to repopulate SOS-deficient intestines, we also assessed the effects of Matrigel alone and conditioned medium used for intestinal organoid cultures. Six days after TMX administration, when SOS1/2 DKO mice already exhibited signs of bacteremia and weight loss, mice were transplanted with 1,000 WT intestinal organoids suspended in 5% Matrigel per mouse, 5% Matrigel in PBS, or 50% L-WRN conditioned medium (CM) in PBS. Notably, all treatments resulted in a significant extension of lifespan (Fig. 6B). Following surgery, the treated mice showed a transient reduction in body weight, followed by progressive recovery and weight gain, while maintaining stable body temperature compared to SOS1/2 DKO mice that did not undergo surgery (Figs. 6C and D).

**Figure 6:**
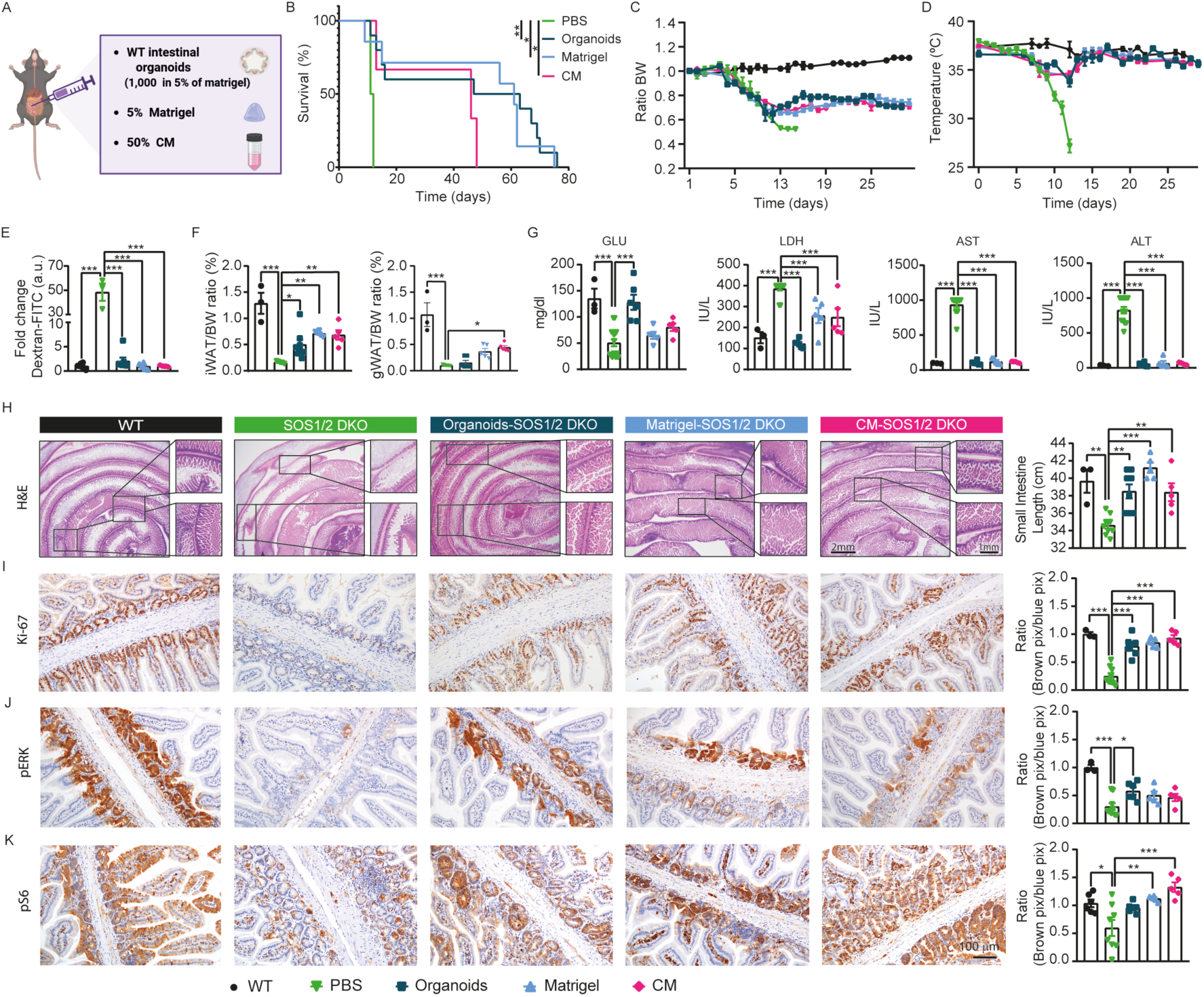
Evaluation of therapeutic intervention strategies in SOS1/2 DKO mice. (A) Scheme of regenerative strategies involving the injection of WT intestinal organoids, Matrigel alone, or L-WRN conditioned medium. *Created with BioRender.com* (B) The survival curves of SOS1/2 DKO mice orthotopically transplanted with organoids, Matrigel, and conditioned media (CM) compared to mice treated with PBS. (C) Body weight normalized with the first day before TMX administration (day 0). (D) Body temperature over time in control and surgically treated mice. (E) Serum FITC-DEAD-dextran 4KDa (FITC-Dextran) levels were evaluated after surgery. (F) Fat pads (iWAT and gWAT) weights were normalized against body weight after 30 days of surgery. (G) Glucose (GLU), Lactate dehydrogenase (LDH), Aspartate aminotransferase (AST), and Alanine aminotransferase (ALT) were measured in the serum of all experimental groups on day 30 after surgery. (H) Representative H&E sections of the small intestine of WT mice, SOS1/2 DKO mice orthotopically implanted with PBS, organoids, Matrigel, and conditioned media (CM), and mice after 30 days of the surgery, together with the small Intestine length quantification. (I-K) Representative immunohistochemistry images showing Ki-67 (I), pERK (J), and pS6 (K) staining in WT, SOS1/2 DKO mice orthotopically implanted with PBS, organoids, Matrigel, and conditioned media (CM) after 30 days of the surgery, with quantification. Data expressed as the mean ± s.e.m. for at least n=5 for each genotype and treatment. Statistical analysis was performed using one-way ANOVA, with Tukey’s post hoc test. Significance indicators: * vs. PBS-treated SOS1/2 DKO mice. *, p<0.05. **, p<0.01; ***, p<0.001.

At 30 days post-surgery, gut permeability was assessed using FITC-Dextran. This time point was chosen to ensure surgical success and to exclude outlier responses. Gut permeability was normalized in all treatments (Fig. 6E). iWAT mass increased across all treatments, although it did not fully reach WT levels, whereas gWAT was significantly increased only in CM-orthotopically implanted mice compared with SOS1/2 DKO animals (Fig. 6F). Importantly, glucose levels and hepatic enzyme markers (AST, ALT, and LDH) returned to values like those observed in WT mice (Fig. 6G).

Histological evaluation (H&E staining) of SOS1/2 DKO mice orthotopically implanted with organoids, Matrigel, or CM revealed marked preservation of intestinal architecture. Treated animals displayed *de novo* crypt formation, enhanced epithelial regeneration, and a significant increase in small intestine length compared to untreated controls, although a subset of crypts remained shortened and focal areas of tissue injury persisted (Fig. 6H). These structural improvements were accompanied by a robust increase in epithelial proliferation, as evidenced by Ki-67 immunostaining, which was restored to levels comparable to WT mice (Fig. 6I). In parallel, pERK staining was selectively enhanced in organoid-implanted mice, albeit not reaching WT levels, whereas S6 phosphorylation was predominantly increased in CM-implanted animals and reached WT levels in all treatments (Fig. 6J and K).

Collectively, these findings indicate that these orthotopic interventions promote intestinal regeneration and preserve epithelial integrity, thereby contributing to the survival of SOS1/2 DKO mice.

## DISCUSSION

We previously reported that SOS1/2 DKO adult mice died suddenly following SOS1 disruption(10). To determine the cause of death, we conducted several analyses. Blood tests revealed clear signs of liver failure, both with oral gavage of TMX and the TMX diet. However, liver failure alone was insufficient to explain the lethality. Similarly, bone marrow transplantation experiments excluded defects in hematopoietic cell populations as a primary cause of death(10).

Consistent with the idea that lethality might arise from systemic failure, we evaluated several therapeutic interventions. Glucose supplementation was tested due to the markedly low glucose levels observed in SOS1/2 DKO mice, along with promising outcomes reported in prior studies(48). However, the modest and transient extension of survival suggests that hypoglycemia is more likely a downstream consequence rather than a primary driver of lethality. Similarly, despite reports of improved outcomes in obese or overweight states, the so-called “obesity paradox”(26), HFD not only failed to provide protection but appeared to exacerbate mortality. Lastly, NAC was tested as a potential mitigator of oxidative stress, given previous observations of elevated oxidative stress in the liver and MEFs derived from DKO mice(15). Unfortunately, antioxidant treatment with NAC did not ameliorate disease severity and instead accelerated death.

Overall, our studies of the consequences of SOS1 and SOS2 ablation in the intestine revealed that the simultaneous loss of both proteins resulted in shortening of the small intestine, reduction in stem cell numbers, decreased proliferation, and reduced ERK and S6 phosphorylation, accompanied by increased intestinal permeability and subsequent sepsis, ultimately leading to death of the mice (Fig. 7).

**Figure 7:**
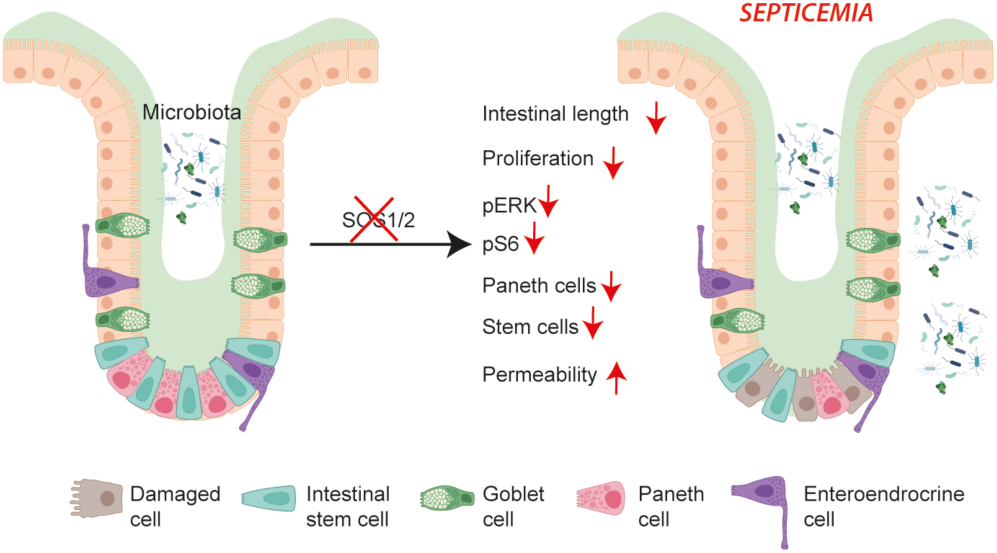
Schematic representation of *in vivo* consequences of simultaneous SOS1 and SOS2 genetic ablation in the mouse intestine. The loss of both SOS1 and SOS2 GEFs resulted in shortening of the small intestine, reduced ISC numbers, decreased proliferation, reduced ERK and S6 phosphorylation, and increased intestinal permeability. Together, these factors lead to severe septicemia, resulting in the death of the mice. *Created with BioRender.com*.

Mechanistically, our findings identify the small intestine as a critical determinant of survival in SOS1/2 DKO mice. Loss of SOS1/2 resulted in a pronounced reduction in ERK phosphorylation and a marked impairment in intestinal epithelial proliferation, highlighting the essential role of SOS-dependent RAS-MAPK signaling in sustaining intestinal epithelial renewal and tissue homeostasis. This interpretation is consistent with previous studies demonstrating that ERK depletion during embryonic development or in adult mice leads to severe intestinal abnormalities, including epithelial disorganization, reduced proliferative capacity, and shortened lifespan(27,28). Interestingly, although SOS1 KO mice also showed reduced ERK phosphorylation, their proliferative capacity remained relatively preserved. This suggests the presence of compensatory mechanisms, likely mediated by SOS2 or alternative signaling pathways, which help maintain epithelial turnover in the absence of SOS1 alone.

The maintenance of intestinal homeostasis requires both sustained proliferation and the coordinated presence of distinct cell types, such as Paneth and ISCs. Paneth cells play crucial roles in protecting against microbial invasion, regulating stem cell dynamics, and modulating immune responses in the gut(49), and sustain LGR5⁺ ISCs by providing key niche factors, including EGF, Wnt, and Notch ligands(50). Additionally, suppressing MAPK activity in the crypts enhances Wnt/β-catenin signaling, promoting Paneth cell differentiation(25). In line with this, SOS1 KO mice exhibited an increase in Paneth cells, consistent with reduced ERK phosphorylation, whereas SOS1/2 DKO mice did not show this increase and instead displayed a significant depletion of ISCs (LGR5⁺/CD44⁺), indicating that the combined loss of both SOS proteins critically impairs the ISC niche and compromises regenerative capacity and barrier maintenance. Remarkably, despite upregulation of Wnt pathway-related genes in SOS1/2 DKO mice, the expression of genes essential for ISC maintenance was insufficient to preserve crypt architecture(51,52). While other epithelial and stromal populations can partially compensate for Paneth loss and support ISC maintenance(53), this compensatory plasticity has limits. Importantly, beyond cellular plasticity, controlling inflammation and its downstream consequences is crucial for maintaining epithelial integrity. Chronic or unresolved inflammation induces profound metabolic stress, particularly at the mitochondrial level, impairing ISC renewal and driving their transition into dysfunctional Paneth-like cells, as shown in Crohn’s disease models(54).

It is worth noting that TMX administration itself can impair ISC maintenance by enhancing fatty acid degradation and causing mitochondrial damage, thereby compromising epithelial regeneration(55). However, despite identical TMX exposure across all genotypes, only SOS1/2 DKO mice failed to recover, developing progressive epithelial barrier dysfunction characterized by increased gut permeability and systemic bacterial dissemination, establishing barrier failure as the primary driver of sepsis and lethality. A key finding supporting this conclusion is that antibiotic treatment, while partially improving survival, failed to restore barrier integrity, confirming that microbial invasion is a consequence of the epithelial failure, not its initial cause. This aligns with recent reports showing that even after effective bacterial clearance, residual microbial products can perpetuate systemic inflammation and contribute to sepsis-related mortality, supporting the notion that epithelial barrier repair is crucial for full recovery(56).

The bacterial dissemination in SOS1/2 DKO mice triggered a systemic inflammatory response characteristic of sepsis. Initially, we observed a hyperinflammatory phase, characterized by elevated circulating neutrophils and monocytes, which was quickly followed by a sharp decline in lymphocyte counts, indicative of immune exhaustion. This biphasic pattern resembles the well-described transition from hyperinflammation to immunosuppression, in which lymphocyte apoptosis and monocyte/macrophage dysfunction increase susceptibility to secondary infections(57). Notably, our model also captured organ dysfunction, as evidenced by increased serum AST and ALT levels coinciding with rising sepsis scores. While earlier reviews focus primarily on cellular and molecular immunosuppressive mechanisms(58), our data provide a temporal link between immune dysregulation and liver injury in SOS1/2 DKO mice.

Importantly, treatment of SOS1/2 DKO mice with intestinal organoids, Matrigel, or conditioned media significantly extended survival. These interventions increased proliferation, ERK, and S6 phosphorylation, indicating that stemness-promoting factors can partially restore proliferative and signaling capacity, thereby counteracting, at least transiently, the rapid lethality of the SOS1/2 DKO phenotype. Each of these treatments is likely to act through distinct but complementary mechanisms. Thus, Matrigel may provide a three-dimensional extracellular matrix that mimics aspects of the native ISC niche, supporting ISC survival and proliferation through enhanced cell-matrix interactions(59). Intestinal organoid transplantation could supply functional ISCs able to integrate into the damaged epithelium and support partial restoration of barrier integrity(44). Conditioned media, which contain soluble niche-derived factors such as Wnt3 and R-spondin, may activate Wnt/β-catenin signaling and transiently support ISC function and differentiation (60,61). Comparable rescue strategies have been reported in other contexts. For example, GSK3β inhibitors were shown to restore Paneth cell numbers and reduce mortality caused by DSS-induced inflammatory bowel disease in Lgr4-deficient mice (52). Moreover, R-spondin, which promotes self-repair and protects against TNF-induced apoptosis (62), has been shown to restore Lgr5^high^ ISC proliferation in URI^(Δ/Δ)Int-Lgr5-EGFP^ mice (63). Consistently, Zhang *et al.* identified lymphatic endothelial cells as a key source of R-spondin 3 in the small intestine, where lymphangiocrine Rspo3 signaling is essential for ISC recovery and epithelial regeneration following cytotoxic injury (64).

Collectively, these findings suggest that the therapeutic benefit observed across these approaches is largely driven by paracrine support, in which the administered matrices, organoids, or soluble niche-derived factors act as a transient source of trophic cues that stimulate endogenous stem cell recovery rather than achieving full structural replacement of the damaged epithelium. Notably, although survival was markedly improved, mortality was not completely prevented, arguing against complete and permanent epithelial replacement as the primary mechanism of action. In this context, our findings (Fig. 7) underscore the indispensable role of SOS1 and SOS2 in maintaining intestinal integrity. Furthermore, beyond defining the mechanisms underlying severe epithelial and metabolic dysfunction, they also highlight potential therapeutic avenues against barrier compromise and systemic sepsis.

## ACKNOWLEDGEMENTS

The authors wish to thank the Pathology Unit, the Mouse Model Experimentation Unit, and the Advanced Cellular Analysis Unit at CIC for their assistance in carrying out this work.

## DATA AVAILABILITY STATEMENT

All data generated or analyzed during this study are included in this published article and its supplementary information files.

## AUTHOR CONTRIBUTIONS

ES, AOS, and RGN contributed to the conceptualization, design, and supervision of the study AOS, RGN, PRR, ADA, CG, NC, RFM, RN, and DD contributed to the experiments, data collection, and analysis. AOS performed data visualization. ES and AFM provided financial support. AOS, ES, and RGN wrote the manuscript, and all authors revised and approved the final version.

## ETHICS DECLARATIONS

All mouse experiments were approved by the Bioethics Committee of the University of Salamanca (#987, #1236) and conducted in its NUCLEUS animal facility according to European (2007/526/CE) and Spanish (RD1201/2005 and RD53/2013) guidelines for animal care and experimentation.

## SUPPLEMENTARY FIGURES

**Figure S1:**
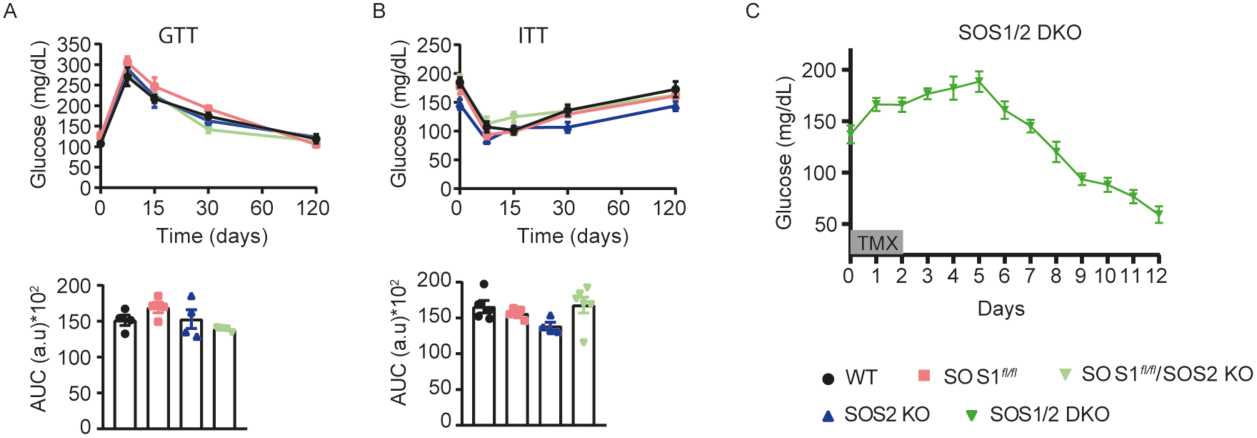
Glucose and insulin tolerance tests reveal no genotype-dependent differences before TMX treatment, whereas SOS1/2 DKO mice exhibit reduced glucose levels following TMX administration. (A) A glucose tolerance test (GTT) was performed before TMX treatment in the indicated genotypes. The upper panel shows blood glucose levels over time, and the lower panel shows quantification of the area under the curve (AUC). (B) An insulin tolerance test (ITT) was performed before TMX treatment in the indicated genotypes. The upper panel shows blood glucose levels over time, and the lower panel shows quantification of the area under the curve (AUC). (C) Serum glucose was measured during TMX treatment on SOS1/2 DKO mice. Data expressed as the mean ± s.e.m. for at least n=4 for each genotype per day.

**Figure S2:**
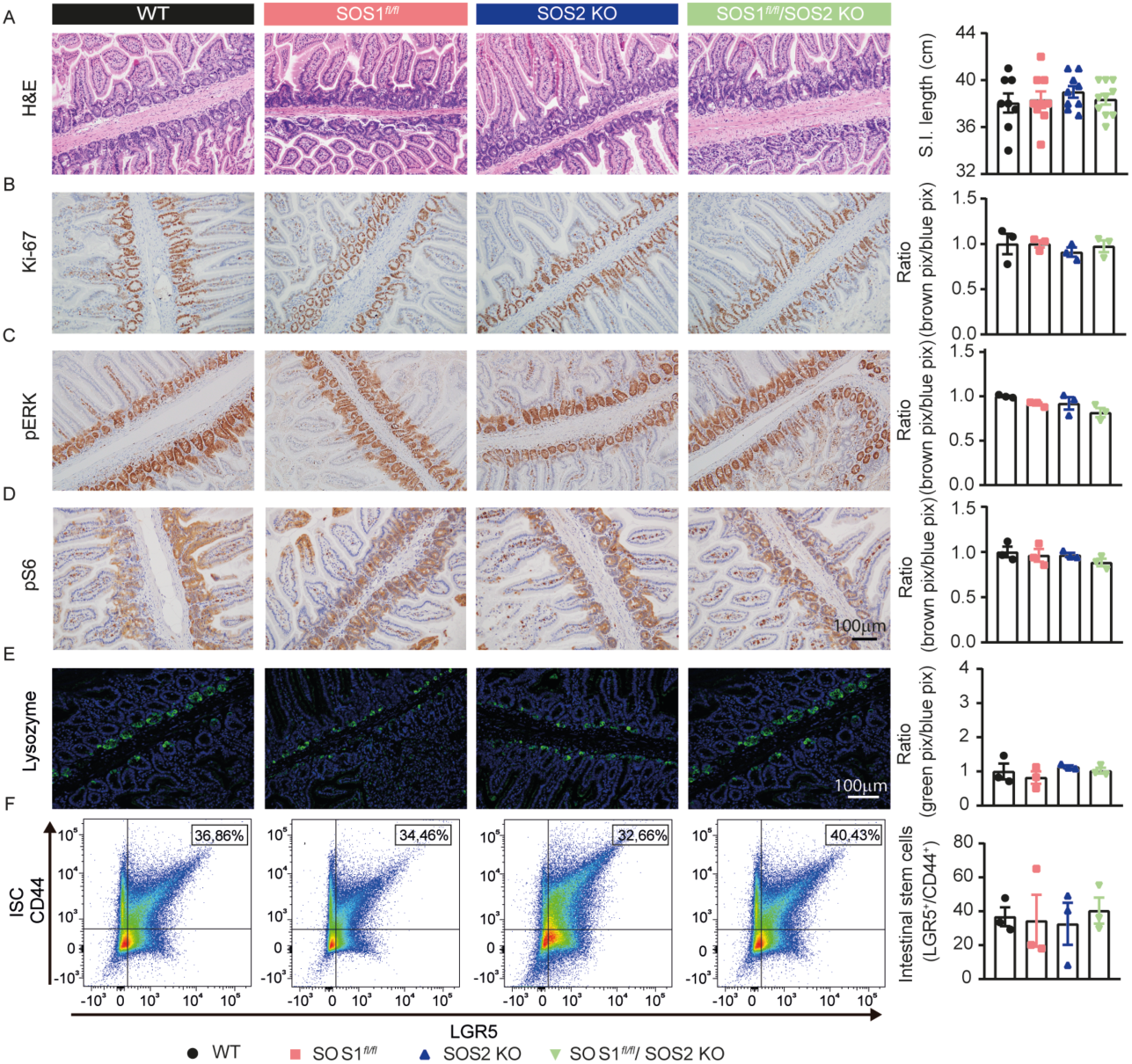
Characterization of the small intestine reveals no genotype-dependent differences before TMX treatment. (A) Representative hematoxylin and eosin (H&E)-stained sections of the small intestine from all experimental groups before TMX treatment, and Small Intestine Length quantification from each experimental group before TMX treatment. (B-D) Representative immunohistochemistry images showing Ki-67 (B), pERK (C), and pS6 (D) staining in WT, SOS1 KO, SOS2 KO, and SOS1/2 DKO mice before TMX treatment, with quantification. (E) Representative lysozyme immunofluorescence images of the small intestine of the four genotypes under study before TMX treatment, with quantification. (F) Representative FACS profiles of LGR5^+^/CD44^+^ expression and the frequency of LGR5^+^/CD44^+^ Intestinal Stem Cells (ISC) quantification from each experimental group before TMX treatment. Data expressed as the mean ± s.e.m. for at least n=4 for each genotype.

**Figure S3:**
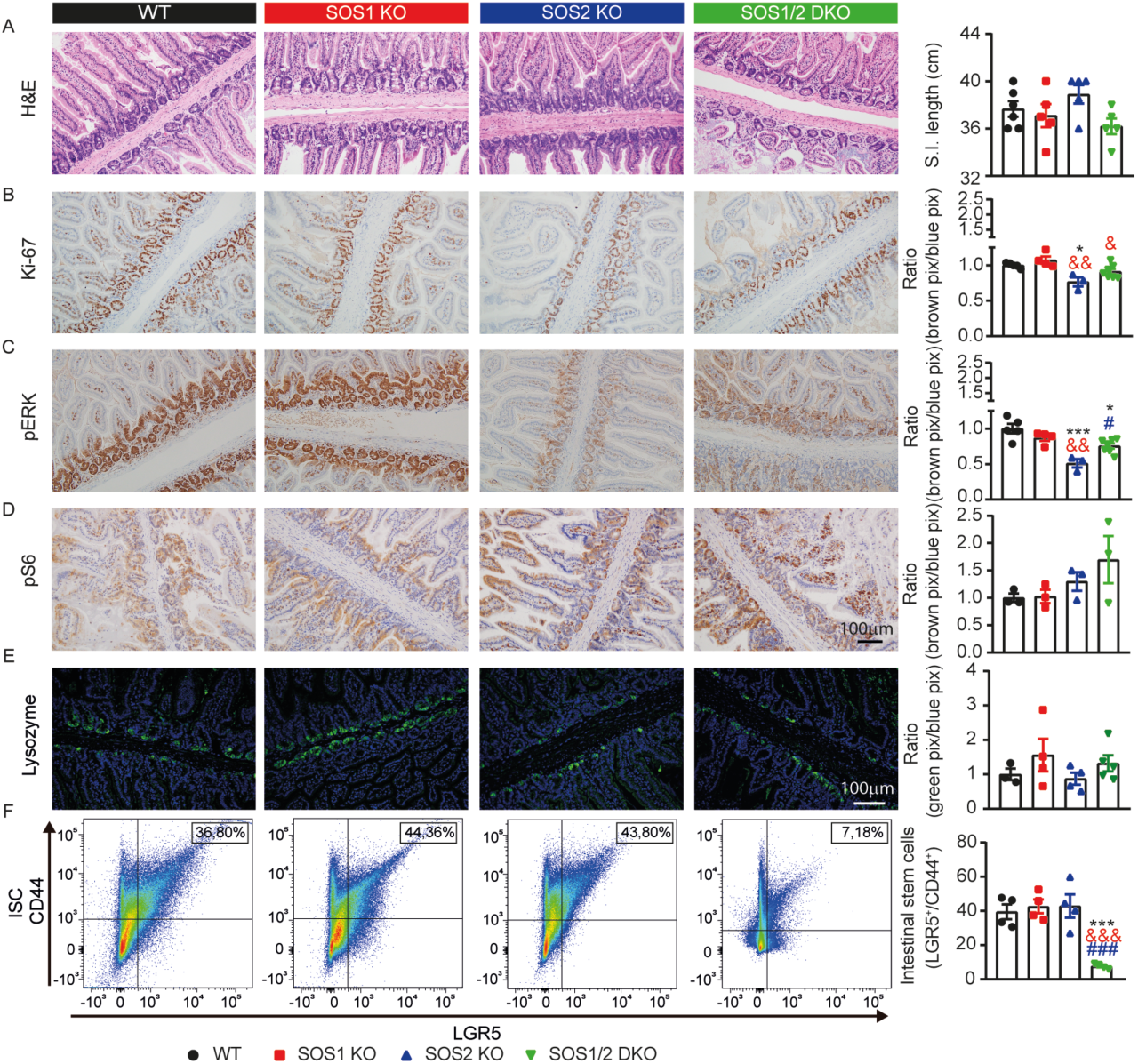
Ki-67 staining and ERK phosphorylation were reduced in SOS2 KO and SOS1/2 DKO mice after 6 days of TMX treatment. (A) Representative hematoxylin and eosin (H&E)-stained sections of the small intestine from all experimental groups on day 6 after TMX treatment, and Small Intestine Length quantification from each experimental group on day 6 after TMX. (B-D) Representative immunohistochemistry images showing Ki-67 (B), pERK (C), and pS6 (D) staining in WT, SOS1 KO, SOS2 KO, and SOS1/2 DKO mice 6 days after TMX treatment, with quantification. (E) Representative lysozyme immunofluorescence images of the small intestine of the four genotypes under study on day 6 after TMX treatment, with quantification. (F) Representative FACS proles of LGR5^+^/CD44^+^ expression and the frequency of LGR5^+^/CD44^+^ Intestinal Stem Cells (ISC) quantification from each experimental group on day 6 after TMX treatment. Data expressed as the mean ± s.e.m. for at least n=4 for each genotype. Statistical analysis was performed using one-way ANOVA, with Tukey’s post hoc test. Significance indicators: * vs. WT, & vs. SOS1 KO and # vs. SOS2 KO; *,& p<0.05; && p<0.01; *** p<0.001.

**Figure S4:**
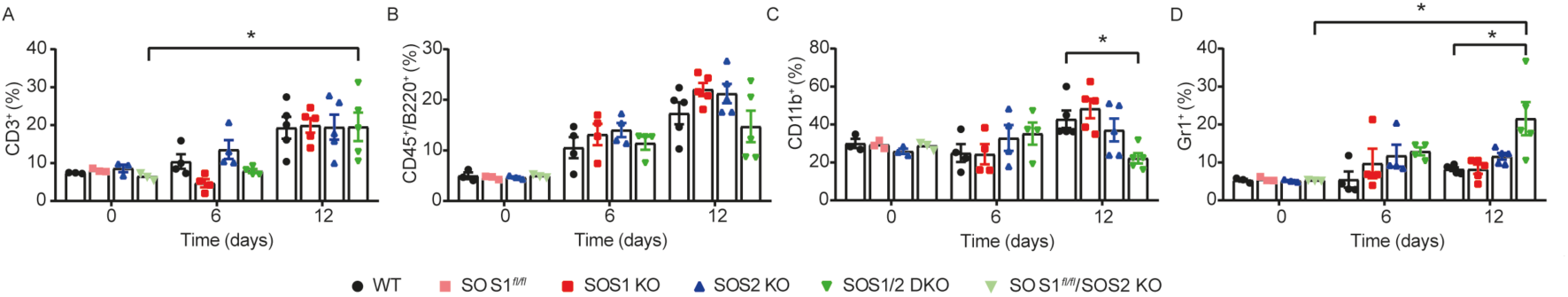
Dynamic alterations in lymphoid and myeloid cell populations following experimental induction. (A-D) The percentage of CD3^+^, CD45^+^/B220^+^, CD11b^+^, and Gr1^+^ cells in peritoneal lavage was assessed on days 0, 6, and 12 of TMX treatment in all genotypes. Data expressed as the mean ± s.e.m. for at least n=3 for each genotype per day. Statistical analysis was performed using one-way ANOVA, with Tukey’s post hoc test. Significance indicators: * vs. WT, & vs. SOS1 KO, and # vs. SOS2 KO; * p<0.05.

**Table S1:**
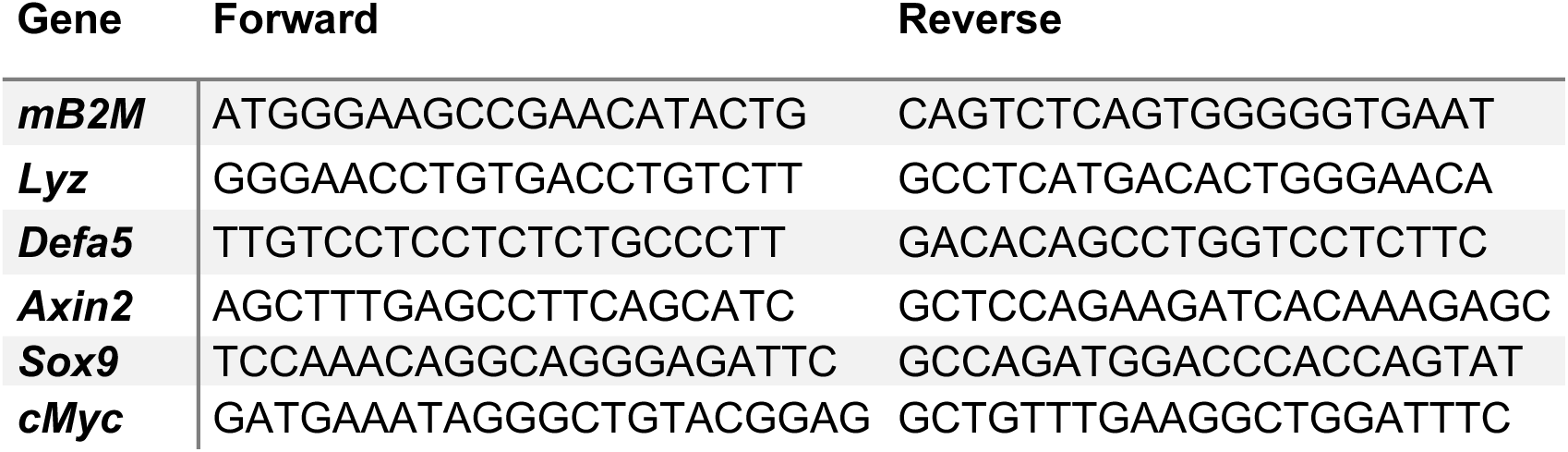
Primer sequences used for qRT-PCR.

**Table S2:**
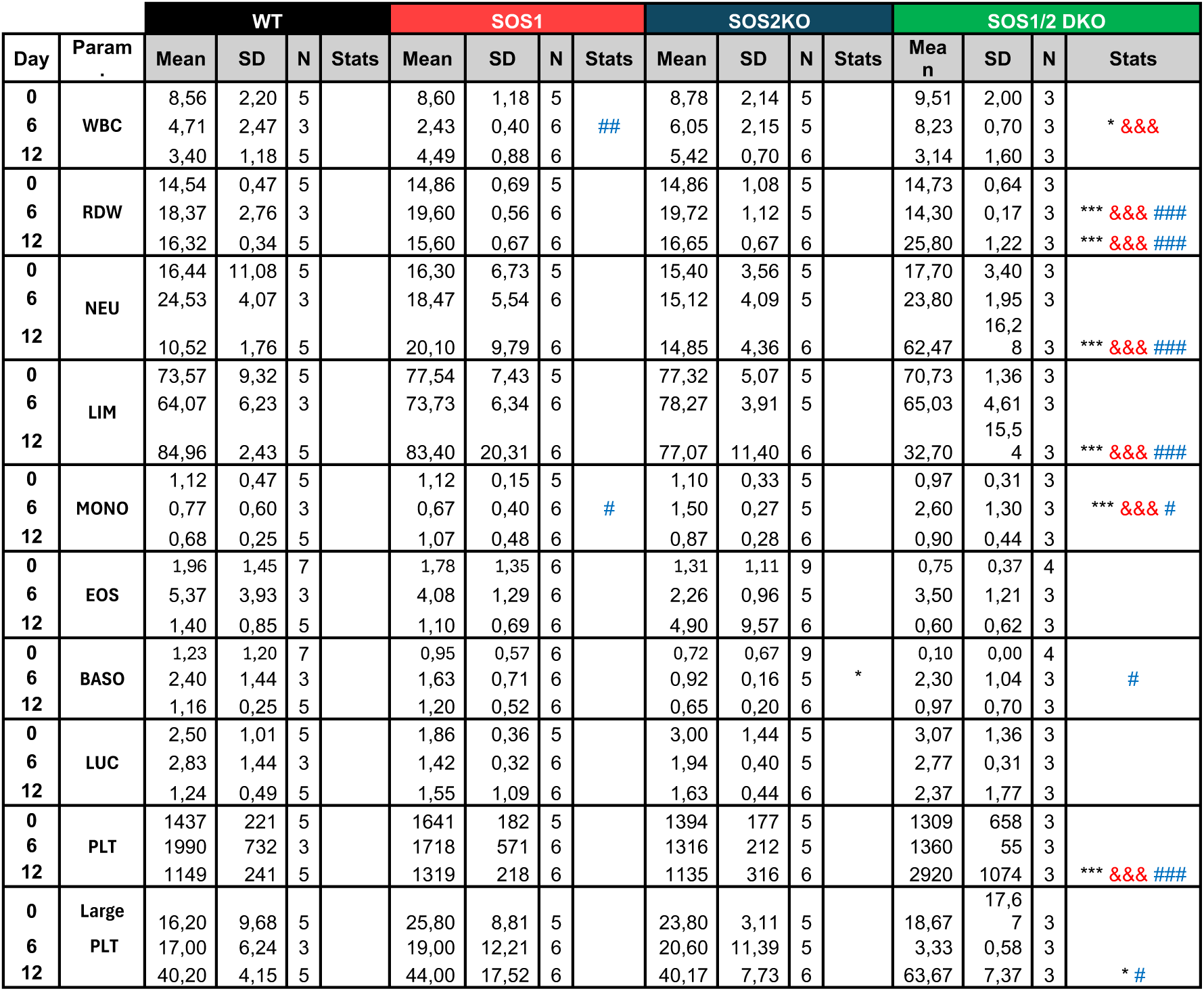
Progressive changes in haematological parameters following experimental induction. Immune profiling of WT, SOS1 KO, SOS2 KO, and SOS1/2 DKO at days 0, 6, and 12 during TMX treatment. Data expressed as the mean ± s.e.m. for at least n=3 for each genotype per day. Statistical analysis was performed using one-way ANOVA, with Tukey’s post hoc test. Significance indicators: * vs. WT, & vs. SOS1 KO, and # vs. SOS2 KO; *, &, # p<0.05; ## p<0.01; ***, &&&, ### p<0.001.

